# coronaSPAdes: from biosynthetic gene clusters to RNA viral assemblies

**DOI:** 10.1101/2020.07.28.224584

**Authors:** Dmitry Meleshko, Iman Hajirasouliha, Anton Korobeynikov

**Affiliations:** Tri-Institutional PhD Program in Computational Biology and Medicine, Weill Cornell Medical College, 10021, New York, USA; Center for Algorithmic Biotechnology, St. Petersburg State University, 199004, St. Peterburg, Russia; Institute for Computational Biomedicine, Department of Physiology and Biophysics, Weill Cornell Medicine of Cornell University, NY, 10021, USA; Englander Institute for Precision Medicine, The Meyer Cancer Center, Weill Cornell Medicine, NY, 10021, USA; Department of Statistical Modelling, St. Petersburg State University, 198504, St. Peterburg, Russia

## Abstract

**Motivation:** The COVID-19 pandemic has ignited a broad scientific interest in viral research in general and coronavirus research in particular. The identification and characterization of viral species in natural reservoirs typically involves *de novo* assembly. However, existing genome, metagenome and transcriptome assemblers often are not able to assemble many viruses (including coronaviruses) into a single contig. Coverage variation between datasets and within dataset, presence of close strains, splice variants and contamination set a high bar for assemblers to process viral datasets with diverse properties.

**Results:** We developed coronaSPAdes, a novel assembler for RNA viral species recovery in general and coronaviruses in particular. coronaSPAdes leverages the knowledge about viral genome structures to improve assembly extending ideas initially implemented in biosyntheticSPAdes. We have shown that coronaSPAdes outperforms existing SPAdes modes and other popular short-read metagenome and viral assemblers in the recovery of full-length RNA viral genomes.

**Availability:** coronaSPAdes version used in this article is a part of SPAdes 3.15 release and is freely available at http://cab.spbu.ru/software/spades.

**Supplementary information:** Supplementary data are available at *Bioinformatics*

## 1 Introduction

The COVID-19 pandemic has increased a scientific interest in coronavirus research. The analysis of the coronavirus dataset starts with obtaining full-length virus genome sequence that can be performed using read alignment (Sah *et al*., 2020; Yin, 2020) or de novo assembly (Sah *et al*., 2020; Kim *et al*., 2020).

The assembly pipeline based on read alignment is a tool of choice for the same strains of the close species, e.g. for SARS-CoV-2 SNP profiling of confirmed COVID-19 patients. De novo assembly is better suited for novel species recovery since read alignment for distant species is unreliable. Recently, there were multiple studies that used MEGAHIT (Li *et al*., 2015) assembler to recover full-length sequence of the SARS-CoV-2 genome, also previous studies show that different SPAdes (Nurk *et al*., 2013) modes perform well in virus recovery (Sutton *et al*., 2019; Roux *et al*., 2017) from complex metagenomes and metaviromes. Nevertheless, none of these assemblers was initially designed for viral assemblies: MEGAHIT and metaSPAdes (Nurk *et al*., 2017) are metagenomic assemblers, SPAdes (Prjibelski *et al*., 2020) is designed to assemble single-cell and isolate bacterial datasets, rnaSPAdes (Bushmanova *et al*., 2019) is intended for accurate isoform separation from eukaryotic data. For RNA viral samples (metaviromes and metatranscriptomes) these assemblers can produce fragmented assemblies due to specific sequencing artifacts, coverage variations, host contamination, multiple strains and quasispecies, and splice events (Viehweger *et al*., 2019). Fast and correct assembly and characterization of viral species is a key step in predicting and preventing the future outbreaks.

Many RNA viruses (including coronaviruses) have a conserved gene structure (Masters, 2006; Harrach, 2014; Dadonaite *et al*., 2019; Watts *et al*., 2009) that can help to better assemble full-length genomes. In this study, we present coronaSPAdes — a novel assembler designed for RNA viral data. coronaSPAdes extends ideas of biosyntheticSPAdes, treating conserved segments of the viral genome as domains of biosynthetic gene clusters. While coronaSPAdes was initially developed having coronaviral species in mind, we demonstrate that overall approach is generic and applicable to assembly of other broad viral families.

We show that coronaSPAdes is able to recover full-length genomes from publicly available datasets where other popular assemblers produce fragmented assembly.

## 2 Methods

RNA virus assemblers (Hunt *et al*., 2015; Ruby *et al*., 2013; Yang *et al*., 2012; Baaijens *et al*., 2017) have to face a number of challenges in order to assemble the sequence data into a consensus sequence. These challenges stem from the nature of the sequencing data due to the biases in the reverse transcription and polymerase chain reaction amplification process that current sequencing methods rely on. These biases are further aggravated by enormous viral population diversity causing lots of SNPs as well as structural variations. Such population diversity is explained by high error rates during the replication process of RNA viruses, essentially they occur as quasispecies (i.e, groups of related genotypes) (Denison *et al*., 2011).

These properties of the data cause assembly fragmentation or, even worse, make certain regions disappear from the assembly. Additionally, some RNA viruses (including the species from the order *Nidovirales* that includes coronaviruses) are known to use discontinuous extension of negative strands to produce multiple mRNAs (Sawicki and Sawicki, 1995). Thus, even in case of a single virus species in the sample, assemblers should be able to deal with multiple produced “isoforms”.

Over the years the SPAdes team produced several assembly pipelines aimed for a wide range of sequencing data and tasks. This includes metaSPAdes (Nurk *et al*., 2017) for the assembly of consensus bacterial genomes from metagenomes, rnaSPAdes (Bushmanova *et al*., 2019) centering around the reconstruction of multiple isoforms from eukaryotic data as well as more specialized versions such as metaviralSPAdes (Antipov *et al*., 2020).

None of these pipelines, however, can handle all the challenges that appear during the assembly of RNA viral data:

1. metaSPAdes cannot cope with the properties and sequencing artifacts typical for RNA-Seq data (outlined below). While it expects multiple species in the input data, overall it assumes relatively uniform coverage across a single bacterial genome. When applied to RNA viral data with uneven coverage and extensive variation, the graph simplification procedures of metaSPAdes (namely, the rare strain disconnector, see (Nurk *et al*., 2017) for more details) could confuse the main genome with high-covered variation and therefore fragment the assembly. This phenomenon is especially severe for complex metaviromes (see Results section, assembly of Inluenza and HIV data).
2. While rnaSPAdes certainly can cope with the specifics of RNA-Seq data, its aim is quite the opposite as required for RNA viral assembly. For RNA viral assembly the main task is to remove possible variation due to quasispecies, strain variation and sequencing artifacts. For RNA transcriptome assemble the aim is to preserve as much variation due to multiple isoforms as possible. This is why the graph simplification procedure of rnaSPAdes is quite “gentle” (see (Bushmanova *et al*., 2019) for more details on the graph simplification procedures) which is further compensated by the isoform restoration procedure. However, the assembly graph of a typical RNA viral dataset (especially a metatranscriptomic / metaviromic one) is much more complex as compared to RNA transcriptome one. As a result, many sequencing artifacts, variation and chimeric connections are still left there which might result in fragmented assemblies, if some part is lost by an accident, or mosaic ones if isoform restoration algorithm would incorrectly resolve variation. Also, we could expect that rnaSPAdes as other RNA assemblers would certainly inflate the genome duplication ratio as multiple possible arrangements of variation might appear in the output.
3. metaviralSPAdes pipeline is based on metaSPAdes and uses coverage-based heuristics in order to detect putative DNA virus sequences (cyclic and linear) from assembly graphs, which fails to work on RNA viral data due to uneven coverage. It does not handle RNA-Seq sequencing artifacts as well.

The outlined issues required us to develop a new assembly graph simplification pipeline that is specifically aimed to take into account the specifics of RNA viral data. coronaSPAdes pipeline consists of two main steps: rnaviralSPAdes and HMMPathExtension.

### 2.1 rnaviralSPAdes

rnaviralSPAdes is a standalone assembler on its own that takes a transcriptome, meta-transcriptome, virome or meta-virome dataset on input. rnaviralSPAdes modifies approaches of metaSPAdes and rnaSPAdes in order to assemble RNA viruses on species level.

#### 2.1.1 Removal of low-complexity (poly-A / poly-T) tips and edges and RNA-seq specific chimeric connections

Analysis in (Bushmanova *et al*., 2019) shows that the majority of the chimeric connections in RNA-Seq data are either single-strand chimeric loops or double-strand hairpins. They are detected by analyzing the graph topology rather than nucleotide sequences or coverage.

Another characteristic of RNA-Seq datasets is the large number of low-complexity regions that originate from poly-A tails resulting from polyadenylation at the ends of mRNAs. To avoid chimeric connections and non-informative sequences low-complexity edges are removed from the de Bruijn graph.

Transcriptome and metatranscriptome datasets could be quite large and input reads often contain billions of distinct k-mers. Therefore rnaviralSPAdes implements removal of low-complexity tips and length 1 edges (by default tips shorter than 200 k-mers and having A/T content more than 80% are removed) on both uncondensed and condensed de Bruijn graph. Early removal of large portion of sequencing artifacts before condensing the edges of de Bruijn graph helps to keep the memory consumption low and reduces the running time of further graph cleaning steps as well.

#### 2.1.2 Preventing gap closure by low-complexity overlaps

There are several approaches that helps to assemble the regions of low coverage. Using multiple k-mers in iterative manner is one of them. Another one is the *gap closure* process: paired-end reads are aligned to the tips (and their neighborhoods) of the graph. And if there are enough paired-end reads that span the gap, then the ends of tips are analyzed for a possible overlap that is shorter than k-mer. If there exists an exact overlap that is longer than 10 bp, then the tips are joined into a single edge at this overlap.

It turned out that the majority of such overlaps for RNA data are again low-complexity sequences containing long stretches of “A”s or “T” with few mismatches. Almost all these overlaps are spurious and therefore the produced connection would be chimeric. We modified gap closure algorithm to ignore such overlaps.

#### 2.1.3 Collapsing of quasispecies

rnaviralSPAdes aims for species-level assemblies (as opposed to strain-level assemblies that are certainly infeasible due to high level of variation), therefore the bulge removal procedure was refined to collapse the variation due to quasispecies. Specifically, rnaviralSPAdes collapses long and similar (with respect to the edit distance) parallel edges in the assembly graph. By default, it does so for edges shorter than 1000 and similar to each other by more than 90% nucleotide identity.

#### 2.1.4 Low-abundant strains disconnector

Unfortunately, strain differences are not only manifested as single nucleotide variations and small insertions or deletions. Such variations (especially for complex datasets containing many species) are caused by highly diverged genome regions, rearrangements, large deletions, parallel gene transfer, etc. Therefore the topology of the de Bruijn graph in the neighborhood of such variations is more complex than a few dozens of bulges complicating the strain variation masking procedure.

metaSPAdes includes a dedicated *edge disconnector algorithm* that uses the coverage ratios between adjacent edges in the assembly graph to identify edges with low coverage ratios as those that most likely originate from rare strains. The algorithm then disconnects such edges from high-covered paths. However, there are important exceptions to this approach, for example, cases when low covered edges are connected to repeats with a high copy number. In order to identify such cases, for each edge *e* a high-covered subgraph connected to *e* is constructed. In the case of repetitive region, we expect to find a component with a total sum of edge length being small.

This repeat-preserving heuristics does not work well for RNA viral datasets due to drastically increased coverage variation compared with metagenomic samples leaving many connections intact. Also, we certainly assume viral genomes to have small genome size and not include high-covered repeats (so all repeats must be intra-species).

In rnaviralSPAdes we use the disconnector algorithm without repeat-preserving heuristics and more conservative thresholds with respect to edge coverage ratio to take into account coverage variation.

#### 2.1.5 Generic assembly graph simplification pipeline

Besides the important changes outlined above, rnaviralSPAdes implements the graph simplification approach similar to consensus assembly graph construction pipeline of metaSPAdes.

As a result, the rnaviralSPAdes pipeline alone allows for removal of the majority of RNA sequencing artifacts and collapsing the variation. However, still there might be very ambiguous cases when an assembler could not remove the errors neither using the coverage-based heuristics nor the graph topology. The Ginger dataset outlined in section 3.2 is a good example. Certainly, it might be possible to tune various assembler heuristics to deal with that particular case, however, the solution will unlikely work on other datasets as it will be overly-aggressive in error elimination. As a result, a different approach is necessary. RNA transcriptome assemblers solve the problem with isoform reconstruction step, potentially inflating the genome duplication ratio and producing misassembled contigs (this phenomenon could be seen on the Figure 4). coronaSPAdes instead uses a HMMPathExtension algorithm.

### 2.2 HMMPathExtension

The second step of the coronaSPAdes pipeline, HMMPathExtension, utilizes the information about viral genome organization to distinguish between putative genomic sequences from uncleaned artifacts.

HMMPathExtension is inspired by HMM-based algorithms of biosyntheticSPAdes (Meleshko *et al*., 2019). It takes a set of HMMs and the assembly graph as input. First, HMMPathExtension aligns HMMs to assembly graph and constructs a domain graph (for details of domain graph construction refer to (Meleshko *et al*., 2019)). In order to construct domain graph for an arbitrary set of HMMs, we had to refine the domain graph construction algorithm from (Meleshko *et al*., 2019).

Among the main assumptions of domain graph construction algorithm as in (Meleshko *et al*., 2019) are: 1) profile HMMs matches are longer than the k-mer length that was used during the de Bruijn graph construction and, 2) different profile HMMs matches can not overlap on the de Bruijn graph. None of these hold for viral protein profile HMMs matches.

Profile HMMs that are used in biosyntheticSPAdes represent long domain sequences, that are significantly longer than usual values of k-mer length *k*. In the current version, if profile HMM match is of length *k* or shorter, it matches to the single k-mer on the de Bruijn graph with prefix that has the best-scored match.

biosyntheticSPAdes guarantees that there are no positions on the de Bruijn graph that are matched more than once. This fact arouses from a distinct nature of biosynthetic gene cluster domains. However, given the unprecedented diversity of viral data and building blocks of viral genomes, viral protein HMM hits can overlap, causing initial domain graph construction algorithm to fail. In order to overcome this problem, we greedily select an arbitrary HMM hit and remove all other hits that overlap with it until there are no overlapping hits. This procedure guarantees that strong and weak edges of the domain graph will be correctly added to the graph.

Similar to biosyntheticSPAdes, HMMPathExtension aims to find paths through the domain graph that traverse significant HMM matches in order and translate them to the assembly graph paths. This way the extracted genomic sequence is supported both by the graph topology and the structure of the genome. The only major difference is that biosyntheticSPAdes assumes that different gene clusters do not overlap in the assembly graph, but in case of viruses, multiple virus strains can be easily presented. Unlike biosyntheticSPAdes, HMMPathExtension does not require to thread through all matches in a connected component of the domain graph. It allows to reconstruct multiple virus sequences of the same family. All paths produced by HMMPathExtension algorithm are maximal by inclusion. That means that we stop growing the path only if we can’t add another node on the domain graph to this path on both sides. This algorithm assumes that heavy contamination or sequencing artifacts are not matched by any viral HMMs, and therefore effectively ignored during domain graph traversal stage.

For coronavirus assemblies coronaSPAdes is bundled with the set of HMMs obtained from Pfam SARS-CoV-2 (El-Gebali *et al*., 2018) (despite the name, these HMMs are quite general and represent the profiles of various proteins that belong to coronaviruses as well as more conserved ones like RNA dependent RNA polymerase (RdRp) that is conserved across all RNA viruses (Venkataraman *et al*., 2018)) enriched with a subset of HMMs for coronaviral protein families studied in (Phan *et al*., 2018).

We need to explicitly emphasize that the approach of HMM-guided assembly is not limited to coronaviral genomes. The HMMPathExtension step allows for a custom HMM database specification effectively enabling HMM-guided assemblies of other genomes using their internal structure. In the Results section, we demonstrate how coronaSPAdes can be used to assemble HIV, Influenza and CoV genome sequences from diverse datasets.

For the custom set of HMMs the performance of the assembly would depend on the sensitivity of the chosen protein profile models. They should be universal enough to match the viral species of interest and cover the whole viral genome representing the majority of viral genes. At the same time they must be specific to a particular viral family to disallow spurious matches. Different databases such as Pfam, U-RVDB-prot (Bigot *et al*., 2020) and vFAM (Skewes-Cox *et al*., 2014) could be subset to create a suitable set of profile HMMs.

## 3 Results

We highlight the features of coronaSPAdes using a wide range of publicly available transcriptome and metatranscriptome datasets that include novel and known coronaviral species. We also show how coronaSPAdes could be used for not only coronaviral assemblies by reproducing Influenza and HIV assembly benchmark from (Hunt *et al*., 2015).

### 3.1 SARS-CoV-2 assembly

First assembly of SARS-CoV-2 virus were produced using MEGAHIT and Trinity and published in (Wu *et al*., 2020) and deposited in Genbank under NC_045512.2 accession (reference length 29903 nt). In order to show that coronaSPAdes is able to handle SARS-CoV-2 sequencing data, we reassembled data from this paper (SRA: SRR10971381) using coronaSPAdes and other tools from the SPAdes family tools. coronaSPAdes produced a single 29872 nt contig that ideally aligns to the NC_045512.2 reference genome. This result is similar to Trinity (29874 nt) and MEGAHIT (29801 bp) results. rnaSPAdes was also able to assemble this data in a single contig (29874 nt). rnaviralSPAdes and metaSPAdes were only able to assemble SARS-CoV-2 genome into two contigs. Therefore, coronaSPAdes has the same performance as other state-of-the-art tools on this dataset.

In order to show, that coronaSPAdes is able to work within a wide coverage range we also performed assembly on datasets that were created by downsampling SARS-CoV-2 dataset. Reads were sampled with 0.1, 0.2, …, 0.9 probabilities to construct nine datasets. Both coronaSPAdes and MEGAHIT were able to assemble a complete SARS-CoV-2 genome from all datasets, despite the fact that in the dataset with shallowest coverage, the nucleotide coverage drops to 4 at some genomic regions.

To validate coronaSPAdes performance on dataset that contains several close coronaviral species, we mixed the SARS-CoV-2 dataset with the bat coronaviral dataset recently published in (Zhou *et al*., 2021) (SRA:SRR14381418, the article already used coronaSPAdes for coronaviral assembly). Among the others, this dataset contains the second closest strain to SARS-CoV-2 virus known up to date (RpYN06, 94% nucleotide identity). We assembled this dataset using coronaSPAdes and MEGAHIT. coronaSPAdes assembled three near-complete coronaviral genomes (SARS-CoV-2, RpYN06, and coronavirus similar to the hedgehog coronavirus 1), while MEGAHIT assembled only two genomes (SARS-CoV-2 and coronavirus similar to the hedgehog coronavirus 1).

These results show the utility of coronaSPAdes in SARS-CoV-2 research, and its ability to restore complete coronaviral genomes in presence of close species and different coverage.

### 3.2 Coronaviral assemblies

Fr4nk is a putative novel Alphacoronavirus detected in a metatranscriptome sequencing library from a Peruvian vampire bat (*Desmodus rotundus*, SRA:ERR2756788). The average coverage is 110x with variation from 2x to 1500x in different regions of the genome. This is a new species of Coronavirus based on RdRP, nucleoprotein, membrane protein and replicase 1a, which all classify this virus an Alphacoronavirus outside of all named sub-genera and most similar to a Pedacovirus.

Ginger is a putative novel Alphacoronavirus detected in a transcriptome sequencing library from a Wildcat (*Felis silvestris*, SRA:SRR7287110). The average coverage is 20x with variation from 230x to 6x in different parts of the genome.

PEDV is a known Alphacoronavirus that causes porcine epidemic diarrhea. It was assembled from a transcriptome sequencing library of epithelial cells of pig intestine (*Sus scrofa*, SRA:SRR10829957). The average coverage is 470x with variation from 30x to 8000x in different parts of the genome.

We assembled Fr4nk, Ginger and PEDV using several specialized virus assemblers (IVA, PRICE, SAVAGE), generic metagenome and transcriptome assemblers (MEGAHIT, metaSPAdes, rnaSPAdes, Trinity (Grabherr *et al*., 2011)) and coronaSPAdes. Versions of tools used are listed in Supplementary Section 1, command lines are in Supplementary Section 2, running times and memory consumption are in Supplementary Section 3.

The overview of the results could be found in Table 1. Conventional viral assemblers (IVA and PRICE) are using a seed-and-extend approach, therefore a seed sequence was required. This property greatly reduces their applicability for novel species search. SAVAGE does not require seed sequence, still was unable to produce any assembly despite runtime allowance of one week (also it demands an estimate for the viral genome coverage which might be a challenge for complex datasets with many species present). It seems that conventional viral assemblers are unable to deal with the specifics of large transcriptome and metatranscriptome datasets and therefore their ability to reconstruct novel viral genomes from complex datasets might be limited. Other assemblers (MEGAHIT, Trinity, metaSPAdes, rnaSPAdes) overall have shown acceptable results, however their performance was not uniform, as none of them was able to assemble complete virus genomes out of all 3 datasets. coronaSPAdes was able to produce whole genomes in all cases.

**Table 1.**
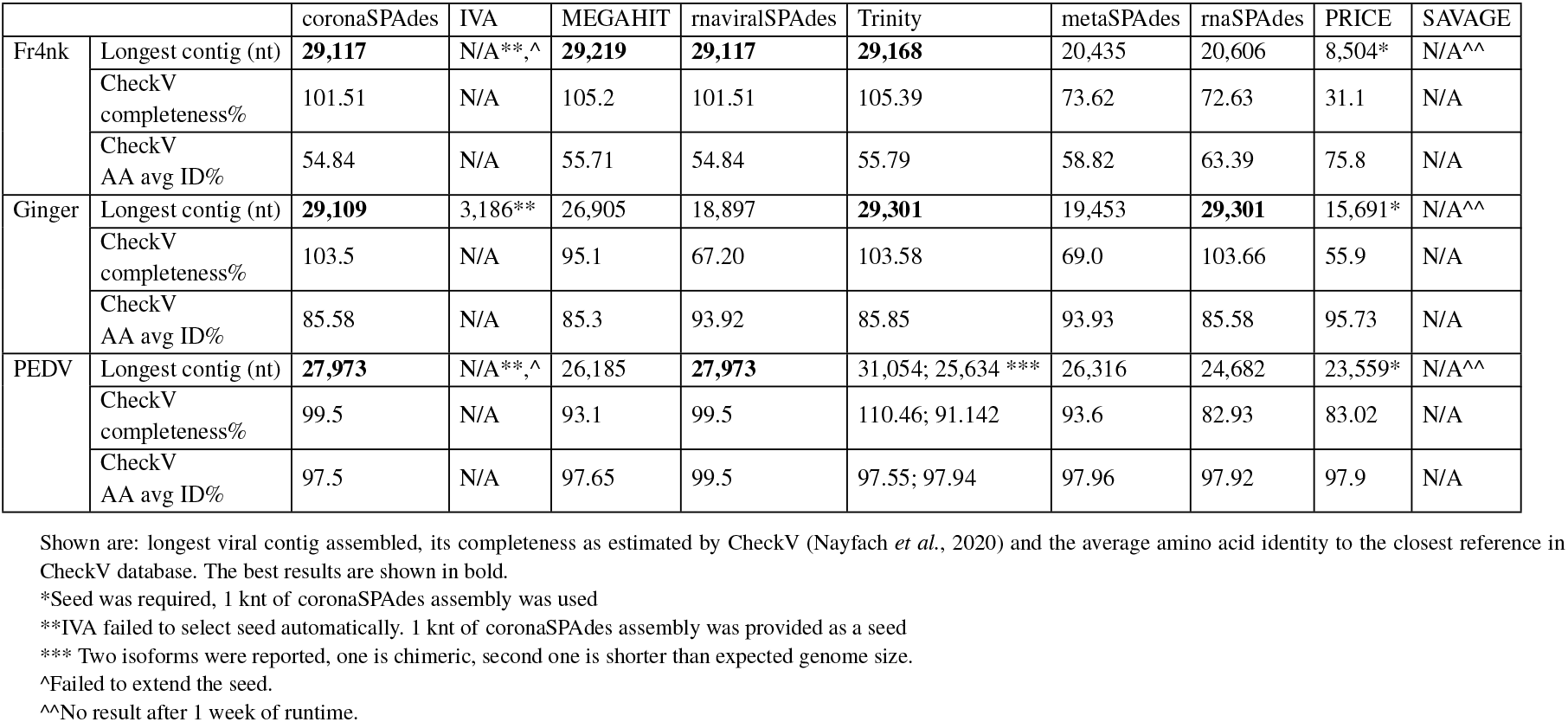
Benchmarking of assemblers on several CoV datasets

### 3.3 HIV and Influenza assemblies

As it was mentioned previously, the HMMPathExtend approach is generic and could be applied to other viral families should the desired set of viral proteins is provided. To showcase this feature we re-create the benchmarking analysis from IVA paper (Hunt *et al*., 2015): we evaluated IVA, Trinity, MEGAHIT, rnaSPAdes, metaSPAdes, rnaviralSPAdes and coronaSPAdes on Illumina paired reads from 68 human immunodeficiency virus 1 (HIV-1) and 172 Influenza A and B virus samples. For this benchmark, coronaSPAdes used the set of HIV and Influenza HMMs extracted from U-RVDB-prot v20 database (Bigot *et al*., 2020).

Figure 2 shows that coronaSPAdes significantly outperforms other tools in terms of assembly contiguity with 30 complete and near-complete assemblies (> 8500 nt). All other assemblers have no more than 14 complete or near-complete assemblies. The number of misassemblies is another important metric that counts the number of times when contig-to-reference alignment has distant breakpoints and shows assembly quality, was calculated using QUAST (Gurevich *et al*. (2013)). coronaSPAdes keeps misassemblies at a relatively low level (see Figure 4), providing a good contiguity-quality trade-off. Also, these results clearly show that metagenomic assemblers might produce suboptimal results and therefore are not suitable for RNA viral assembly from metaviromes.

Influenza assembly is more complicated because influenza type A, B genomes consist of eight segments, that can have highly similar regions at the segment’s ends. As a metric for the assembly contiguity, we sum the longest alignment length across all segments. Figure 3 shows that rnaSPAdes has the best contiguity performance, with coronaSPAdes and Trinity at the second place. However, Figure 4 shows that

MEGAHIT, rnaSPAdes, and Trinity have the worst misassembly statistics (73.0, 19.82, and 7.162 misassemblies per dataset correspondingly), while coronaSPAdes has 2.895 misassemblies per dataset. Therefore coronaSPAdes has a reasonable contiguity-correctness trade-off.

Raw assembly results are available in Supplementary Tables 1 (for Influenza datasets) and 2 (for HIV datasets).

### 3.4 Discovery of novel coronaviruses

coronaSPAdes was used in the Serratus project (Edgar *et al*., 2020) for a widespread search of novel CoV and CoV-like species from public sequencing libraries. From a screen of 3.8 million public RNA-seq, meta-genome, meta-virome and meta-transcriptome datasets deposited in NCBI SRA comprising 5.6 petabases of sequencing reads, 52,772 runs potentially containing CoV sequencing reads were identified. 11,120 of the resulting assemblies contained putative CoV contigs, of which 4,179 aligned to CoV RdRp. These assemblies include sequences from 13 previously uncharacterized or unavailable CoV or CoV-like operational taxonomic units (OTUs), defined by clustering amino sequences of the RdRp gene at 97% identity. 8 of these OTUs were designated to a putative novel genus of coronaviruses, noting that all were found in samples from non-mammal aquatic vertebrates falling outside deltacoronaviruses genus.

The length distribution for assemblies of SRA datasets classified as likely CoV-positive, showing a peak around the typical CoV genome length is presented on Supplementary Figure 1.

## 4 Discussion

Ginger dataset represents the typical case when long sequencing artifacts could influence the assembly results. All metagenome assembles lost some parts of the genome likely being unable to remove the long artifacts having coverage similar to the virus genome (see Figure 1 for assembly graph). RNA assemblers (rnaSPAdes and Trinity) solved this problem via isoform restoration steps: different “isoform” paths across this subgraph were produced and one of them was a full-length viral genome. The downside of this approach is increased genome duplication ratio as multiple paths through the subgraph are produced and in more complex cases some of them might be mosaic. coronaSPAdes traversed all domain matches in order and also produced a full-length viral genome.

**Fig. 1:**
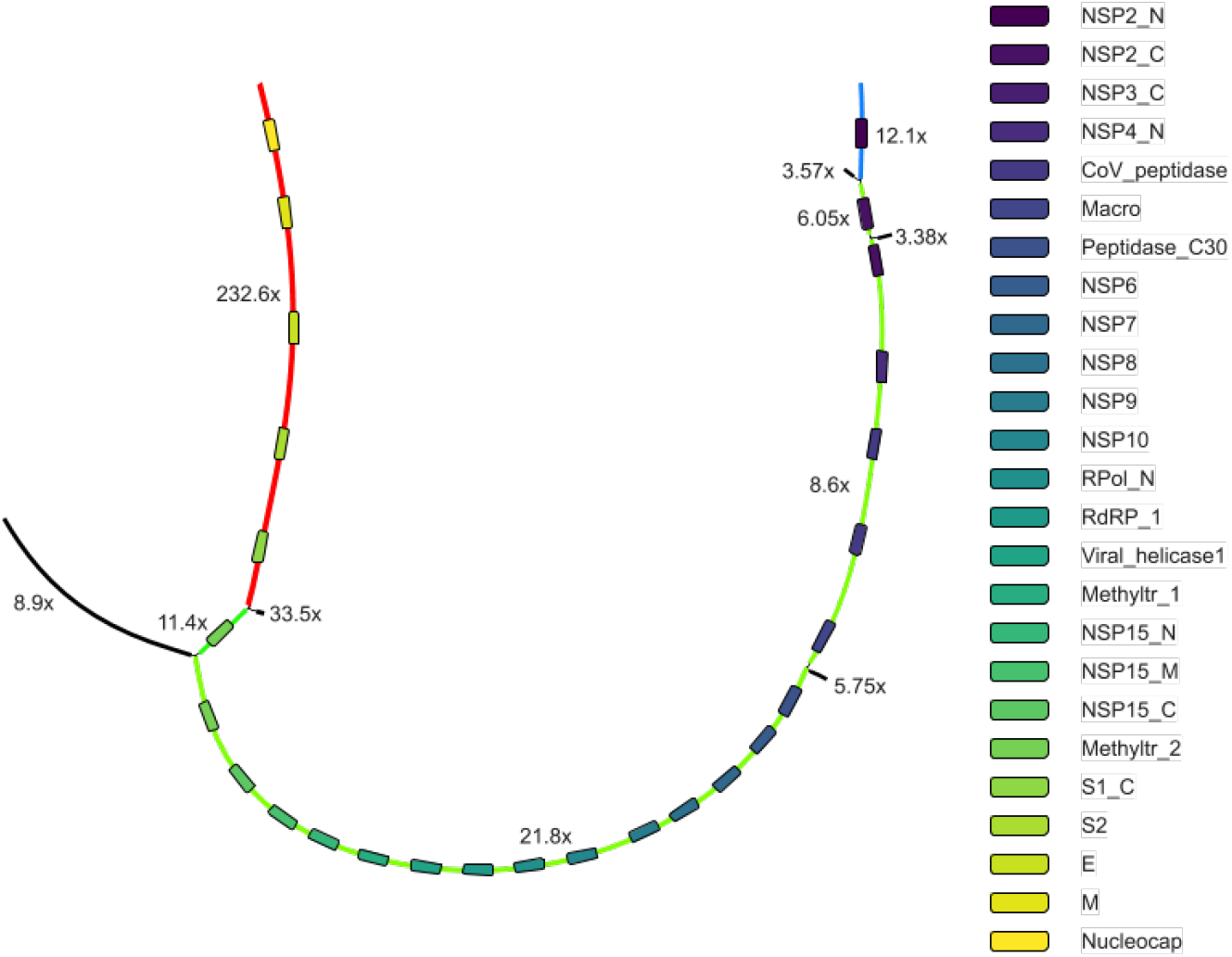
Part of Ginger assembly graph produced with coronaSPAdes. rnaviralSPAdes produced 8 contigs from this subgraph (red, green and blue paths on the graph and 5 black edges), therefore splitting the coronavirus genome into three parts. coronaSPAdes matched viral edges of the graph with domain (rectangles of different color). Path along these matches spells a full-length viral genome

**Fig. 2:**
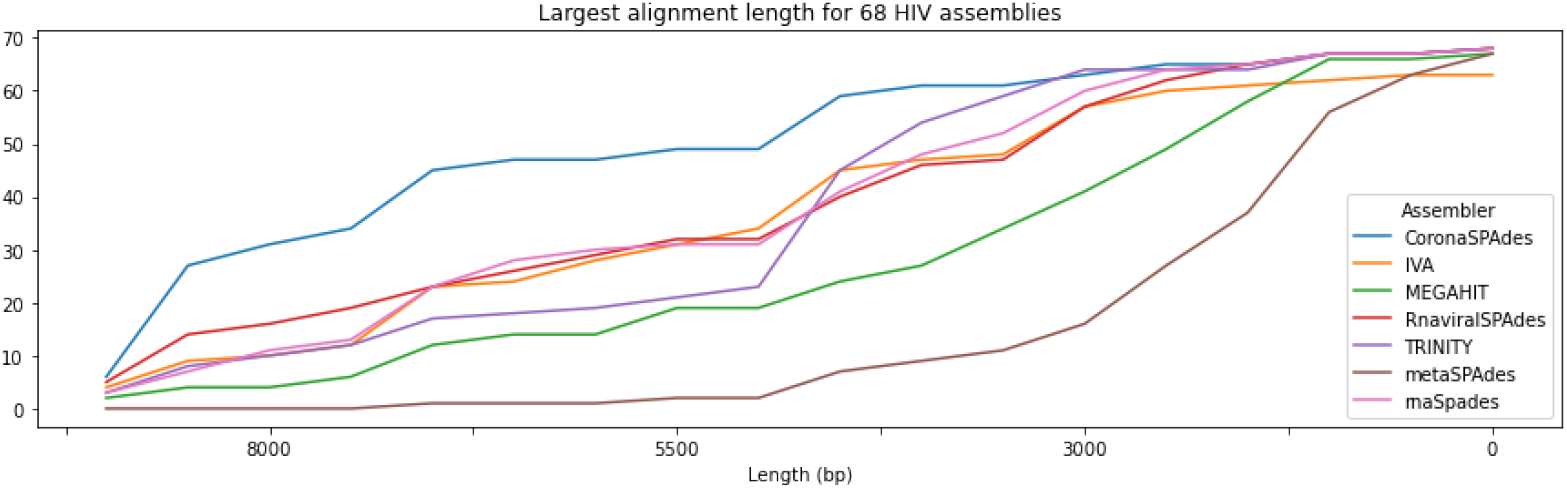
Performance of different assemblers on HIV datasets. Y-axis represents number of datasets which have alignment of such length or greater, similarly to a widely adopted NAx plot.

**Fig. 3:**
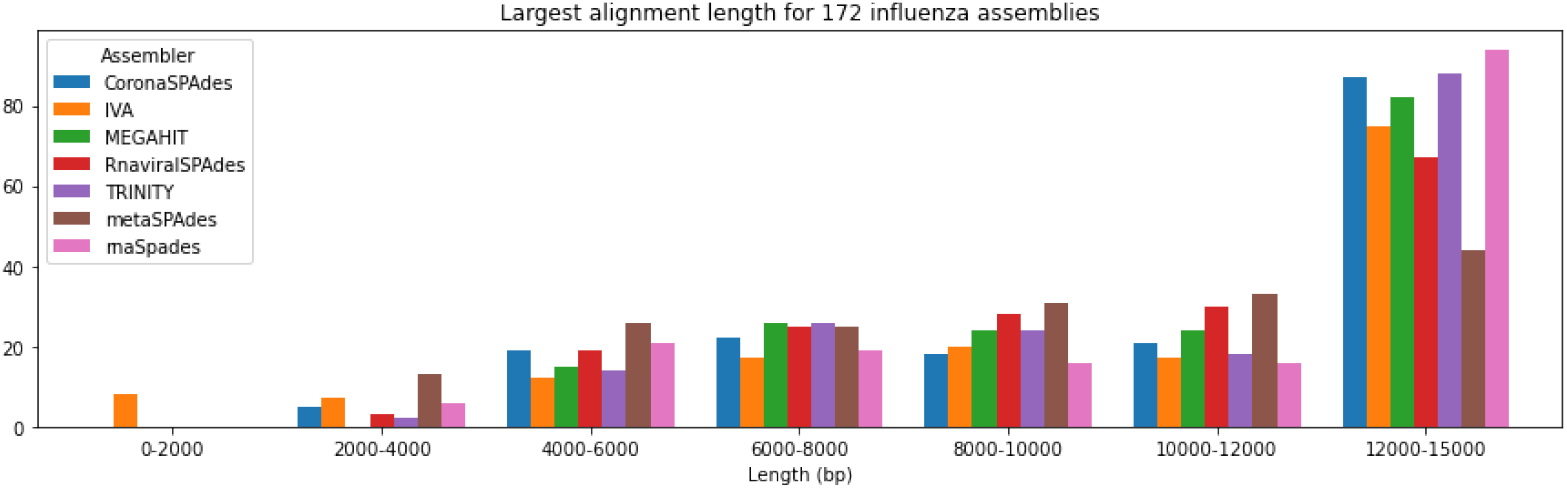
Performance of different assemblers on influenza datasets. Y-axis represents number of datasets which have alignment of such length.

**Fig. 4:**
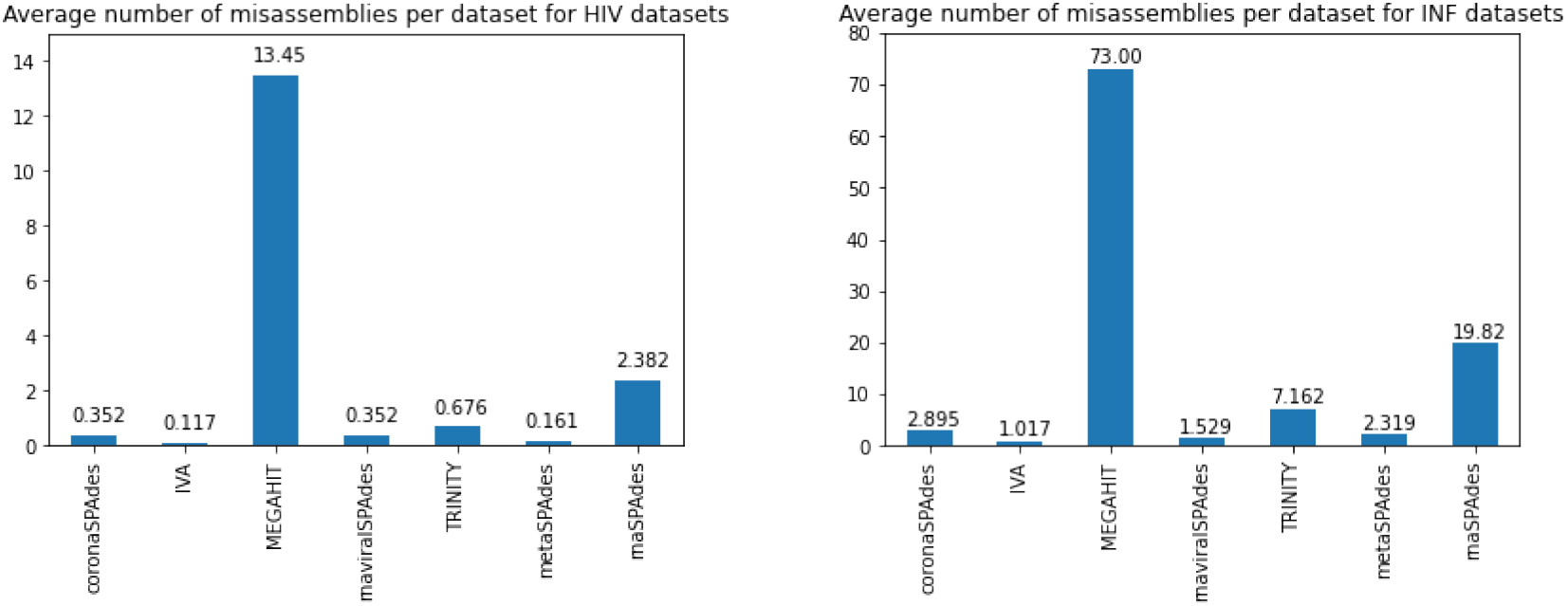
Misassembly statistics for the HIV and influenza datasets. Left: Average number of misassemblies per dataset for HIV datasets. Right: Average number of misassemblies per dataset for influenza datasets.

The influenza A and B virus genomes each comprise eight negative-sense, single-stranded viral RNA segments that code 10–14 proteins, depending on the strain. Each segment possesses noncoding regions, of varying lengths, at both 3’and 5’ ends. However, the extreme ends of all segments are highly conserved among all influenza virus segments (Bouvier and Palese, 2008).

As a result, depending on the particular strain, the segments could appear glued in the assembly graph (see Figure 5), also the Influenza genomes are highly variable with many rearrangements in at least 2 segments due to the antigenic drift and shift processes (Webster and Govorkova, 2014). The challenge for an assembler here is to correctly recover the sequences of all eight segments taking into account possible variation.

**Fig. 5:**
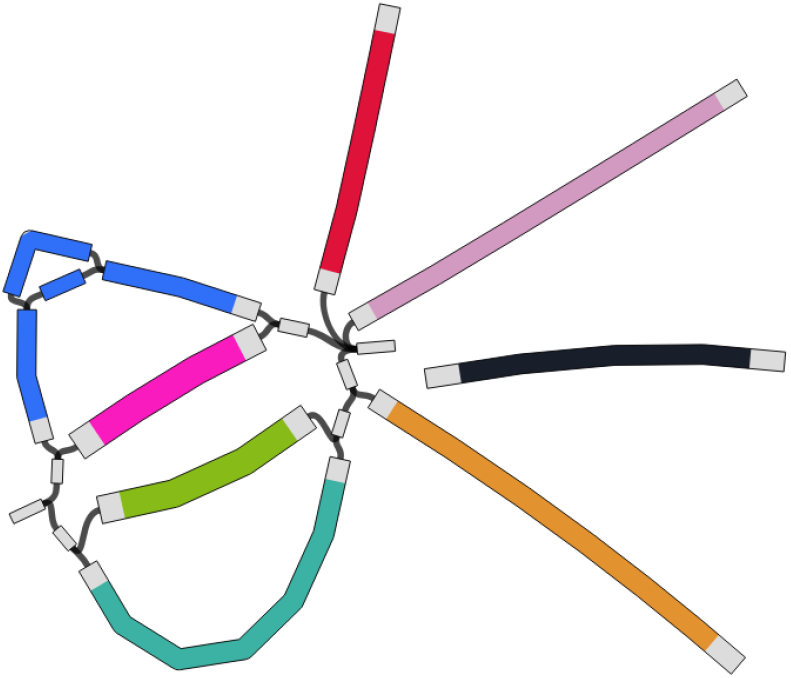
coronaSPAdes assembly graph for ERR732256 Influenza A dataset. Gluing of the genome segments’ terminal parts is typical for influenza assembly graphs. Different CDS in segments are color-coded. There is a variation in HA (blue) segment.

Surprisingly, MEGAHIT for some unknown reason often produced the contigs of 4-8 Knt that contained parts of multiple segments. This results in a highly elevated rate of misassemblies seen there. The results of other assemblers are overall expected as well: both Trinity and rnaSPAdes shown good results in recovery of full-length segments using the isoform reconstruction procedures. However, some segments were clearly mosaic as could be seen from the Figure 4. Seed-and-extend approach of IVA resulted in the most accurate in terms of the average number of misassemblies assemblies, albeit at the expense of the recovery of the fuller segments. coronaSPAdes was able to recover much more still having an acceptable misassembly rate.

HIV genome represents a true nightmare from the assembly standpoint as it employs a very complex system of differential RNA splicing to obtain more than 30 mRNA species from a less than 10 kb genome (Schwartz *et al*., 1990). This results in a very complex and tangled assembly graph (see Figure 6) that an assembler must traverse in order to recover the complete virus genome. Here the HMM-guided approach of coronaSPAdes clearly allows to extend the contigs and recover significantly fuller genomes as compared to other tools.

**Fig. 6:**
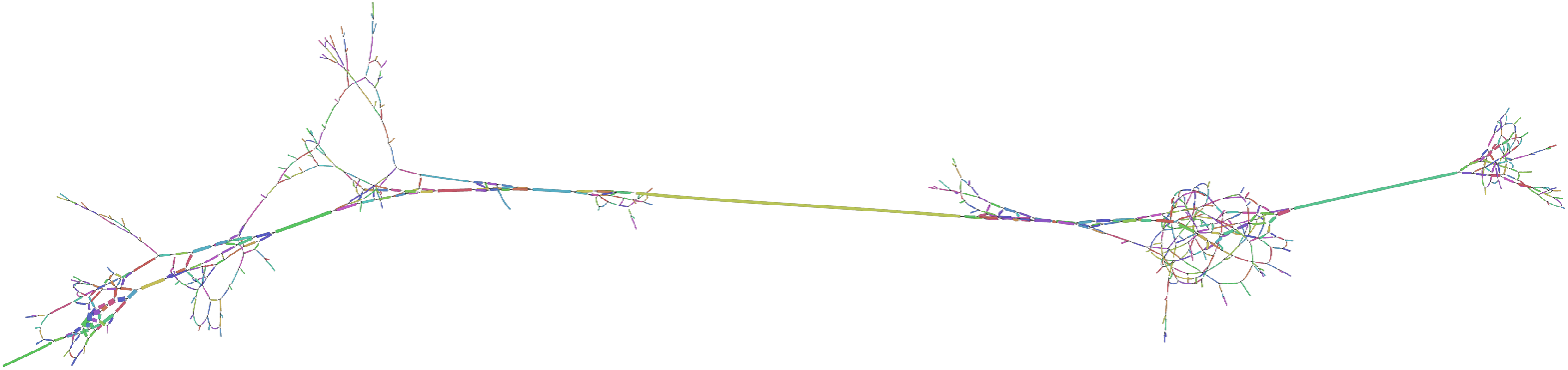
coronaSPAdes assembly graph of a ERR732083 HIV dataset. Tangles in the graph represent high variation in some segments of HIV genome. Graph consists of 856 edges of 130 knt total length with 122 dead-end edges.

## 5 Conclusion

Clearly, assembling RNA viral genomes is very challenging (Hunt *et al*., 2015). The variety of possible kinds of input data and the overall diversity of the species multiplies these challenges even more. We demonstrated that additional information about the genome structure could significantly improve the viral genome recovery even from very complex datasets and therefore catalyze the new viral discoveries.

## 6 Data Availability

The datasets supporting the conclusions of this article are available in the NCBI SRA repository, under accessions SRR10971381 (SARS-CoV-2), SRR14381418 (dataset with RpYN06), ERR2756788 (Fr4nk), SRR7287110 (Ginger), SRR10829957 (PEDV), SRR10971381 (SARS-CoV-2) and SRR14381418 (RpYN06 and other coronaviruses). Access instructions for Serratus data can be found at https://github.com/ababaian/serratus/wiki/Access-Data-Release. Influenza and HIV-1 data were taken from IVA paper (Hunt *et al*., 2015) (lists of accessions are available from Supplementary Tables 1 and 2). coronaSPAdes version used in this article is a part of SPAdes 3.15 release and is available at http://cab.spbu.ru/software/spades and also deposited on Zenodo under doi:10.5281/zenodo.4438269. HIV and Influenza HMMs were extracted from U-RVDB-prot v20 database and are available from http://cab.spbu.ru/software/coronaspades.

## Acknowledgements

The research was carried out in part by computational resources provided by the Resource Center “Computer Center of SPbU”. The authors are grateful to Saint Petersburg State University for the overall support of this work. DM is grateful to T32 Weill Cornell Tri-Institutional Training Program in Computational Biology and Medicine.

## References

Antipov, D., Raiko, M., Lapidus, A., and Pevzner, P. A. (2020). MetaviralSPAdes: assembly of viruses from metagenomic data. Bioinformatics. btaa490.

Baaijens, J. A., Aabidine, A. Z. E., Rivals, E., and Schönhuth, A. (2017). De novo assembly of viral quasispecies using overlap graphs. Genome Research, 27(5), 835–848.

Bigot, T., Temmam, S., Pérot, P., and Eloit, M. (2020). RVDB-prot, a reference viral protein database and its HMM profiles [version 2; peer review: 2 approved]. F1000Research, 8, 530.

Bouvier, N. M. and Palese, P. (2008). The biology of influenza viruses. Vaccine, 26, D49–D53.

Bushmanova, E., Antipov, D., Lapidus, A., and Prjibelski, A. D. (2019). rnaSPAdes: a de novo transcriptome assembler and its application to RNA-Seq data. GigaScience, 8(9). giz100.

Dadonaite, B., Gilbertson, B., Knight, M. L., Trifkovic, S., Rockman, S., Laederach, A., Brown, L. E., Fodor, E., and Bauer, D. L. V. (2019). The structure of the influenza a virus genome. Nature Microbiology, 4(11), 1781–1789.

Denison, M. R., Graham, R. L., Donaldson, E. F., Eckerle, L. D., and Baric, R. S. (2011). Coronaviruses. RNA Biology, 8(2), 270–279.

Edgar, R. C., Taylor, J., Altman, T., Barbera, P., Meleshko, D., Lin, V., Lohr, D., Novakovsky, G., Al-Shayeb, B., Banfield, J. F., Korobeynikov, A., Chikhi, R., and Babaian, A. (2020). Petabase-scale sequence alignment catalyses viral discovery. bioRxiv.

El-Gebali, S., Mistry, J., Bateman, A., Eddy, S. R., Luciani, A., Potter, S. C., Qureshi, M., Richardson, L. J., Salazar, G. A., Smart, A., Sonnhammer, E. L., Hirsh, L., Paladin, L., Piovesan, D., Tosatto, S. C., and Finn, R. D. (2018). The Pfam protein families database in 2019. Nucleic Acids Research, 47(D1), D427–D432.

Grabherr, M. G., Haas, B. J., Yassour, M., Levin, J. Z., Thompson, D. A., Amit, I., Adiconis, X., Fan, L., Raychowdhury, R., Zeng, Q., Chen, Z., Mauceli, E., Hacohen, N., Gnirke, A., Rhind, N., di Palma, F., Birren, B. W., Nusbaum, C., Lindblad-Toh, K., Friedman, N., and Regev, A. (2011). Full-length transcriptome assembly from RNA-seq data without a reference genome. Nature Biotechnology, 29(7), 644–652.

Gurevich, A., Saveliev, V., Vyahhi, N., and Tesler, G. (2013). Quast: quality assessment tool for genome assemblies. Bioinformatics, 29(8), 1072–1075.

Harrach, B. (2014). Adenoviruses: General features. In Reference Module in Biomedical Sciences. Elsevier.

Hunt, M., Gall, A., Ong, S. H., Brener, J., Ferns, B., Goulder, P., Nastouli, E., Keane, J. A., Kellam, P., and Otto, T. D. (2015). IVA: accurate de novo assembly of RNA virus genomes. Bioinformatics, 31(14), 2374–2376.

Kim, J.-M., Chung, Y.-S., Jo, H. J., Lee, N.-J., Kim, M. S., Woo, S. H., Park, S., Kim, J. W., Kim, H. M., and Han, M.-G. (2020). Identification of coronavirus isolated from a patient in korea with COVID-19. Osong Public Health and Research Perspectives, 11(1), 3–7.

Li, D., Liu, C.-M., Luo, R., Sadakane, K., and Lam, T.-W. (2015). MEGAHIT: an ultra-fast single-node solution for large and complex metagenomics assembly via succinct de Bruijn graph. Bioinformatics, 31(10), 1674–1676.

Masters, P. S. (2006). The molecular biology of coronaviruses. In Advances in Virus Research, pages 193–292. Elsevier.

Meleshko, D., Mohimani, H., Tracanna, V., Hajirasouliha, I., Medema, M. H., Korobeynikov, A., and Pevzner, P. A. (2019). Biosyntheticspades: reconstructing biosynthetic gene clusters from assembly graphs. Genome Research, 29(8), 1352–1362.

Nayfach, S., Camargo, A. P., Schulz, F., Eloe-Fadrosh, E., Roux, S., and Kyrpides, N. C. (2020). CheckV assesses the quality and completeness of metagenome-assembled viral genomes. Nature Biotechnology.

Nurk, S., Bankevich, A., Antipov, D., Gurevich, A. A., Korobeynikov, A., Lapidus, A., Prjibelski, A. D., Pyshkin, A., Sirotkin, A., Sirotkin, Y., Stepanauskas, R., Clingenpeel, S. R., Woyke, T., Mclean, J. S., Lasken, R., Tesler, G., Alekseyev, M. A., and Pevzner, P. A. (2013). Assembling single-cell genomes and mini-metagenomes from chimeric mda products. Journal of Computational Biology, 20(10), 714–737. PMID: 24093227.

Nurk, S., Meleshko, D., Korobeynikov, A., and Pevzner, P. A. (2017). metaspades: a new versatile metagenomic assembler. Genome Research, 27(5), 824–834.

Phan, M. V. T., Tri, T. N., Anh, P. H., Baker, S., Kellam, P., and Cotten, M. (2018). Identification and characterization of coronaviridae genomes from vietnamese bats and rats based on conserved protein domains. Virus Evolution, 42).

Prjibelski, A., Antipov, D., Meleshko, D., Lapidus, A., and Korobeynikov, A. (2020). Using SPAdes de novo assembler. Current Protocols in Bioinformatics, 70(1).

Roux, S., Emerson, J. B., Eloe-Fadrosh, E. A., and Sullivan, M. B. (2017). Benchmarking viromics: an in silico evaluation of metagenome-enabled estimates of viral community composition and diversity. PeerJ, 5, e3817.

Ruby, J. G., Bellare, P., and DeRisi, J. L. (2013). PRICE: Software for the targeted assembly of components of (meta) genomic sequence data. G3: Genes, Genomes, Genetics, 3(5), 865–880.

Sah, R., Rodriguez-Morales, A. J., Jha, R., Chu, D. K. W., Gu, H., Peiris, M., Bastola, A., Lal, B. K., Ojha, H. C., Rabaan, A. A., Zambrano, L. I., Costello, A., Morita, K., Pandey, B. D., and Poon, L. L. M. (2020). Complete genome sequence of a 2019 novel coronavirus (sars-cov-2) strain isolated in nepal. Microbiology Resource Announcements, 9(11).

Sawicki, S. G. and Sawicki, D. L. (1995). Coronaviruses use Discontinuous Extension for Synthesis of Subgenome-Length Negative Strands, pages 499–506. Springer US, Boston, MA.

Schwartz, S., Felber, B. K., Benko, D. M., Fenyö, E. M., and Pavlakis, G. N. (1990). Cloning and functional analysis of multiply spliced mRNA species of human immunodeficiency virus type 1. Journal of Virology, 64(6), 2519–2529.

Skewes-Cox, P., Sharpton, T. J., Pollard, K. S., and DeRisi, J. L. (2014). Profile hidden markov models for the detection of viruses within metagenomic sequence data. PLoS ONE, 9(8), e105067.

Sutton, T. D. S., Clooney, A. G., Ryan, F. J., Ross, R. P., and Hill, C. (2019). Choice of assembly software has a critical impact on virome characterisation. Microbiome, 7(1), 12.

Venkataraman, S., Prasad, B., and Selvarajan, R. (2018). RNA dependent RNA polymerases: Insights from structure, function and evolution. Viruses, 10(2), 76.

Viehweger, A., Krautwurst, S., Lamkiewicz, K., Madhugiri, R., Ziebuhr, J., Hölzer, M., and Marz, M. (2019). Direct RNA nanopore sequencing of full-length coronavirus genomes provides novel insights into structural variants and enables modification analysis. Genome Research, 29(9), 1545–1554.

Watts, J. M., Dang, K. K., Gorelick, R. J., Leonard, C. W., Jr, J. W. B., Swanstrom, R., Burch, C. L., and Weeks, K. M. (2009). Architecture and secondary structure of an entire HIV-1 RNA genome. Nature, 460(7256), 711–716.

Webster, R. G. and Govorkova, E. A. (2014). Continuing challenges in influenza. Annals of the New York Academy of Sciences, 1323(1), 115– 139.

Wu, F., Zhao, S., Yu, B., Chen, Y.-M., Wang, W., Song, Z.-G., Hu, Y., Tao, Z.-W., Tian, J.-H., Pei, Y.-Y., et al. (2020). A new coronavirus associated with human respiratory disease in china. Nature, 579(7798), 265–269.

Yang, X., Charlebois, P., Gnerre, S., Coole, M. G., Lennon, N. J., Levin, J. Z., Qu, J., Ryan, E. M., Zody, M. C., and Henn, M. R. (2012). De novo assembly of highly diverse viral populations. BMC Genomics, 13(1), 475.

Yin, C. (2020). Genotyping coronavirus sars-cov-2: methods and implications. Genomics, 112(5), 3588 – 3596.

Zhou, H., Ji, J., Chen, X., Bi, Y., Li, J., Wang, Q., Hu, T., Song, H., Zhao, R., Chen, Y., et al. (2021). Identification of novel bat coronaviruses sheds light on the evolutionary origins of sars-cov-2 and related viruses. Cell.

