## Supplementary Text for "coronaSPAdes: from biosynthetic gene clusters to RNA viral assemblies"

July 20, 2021

### 1 Software versions

1. SPAdes, metaSPAdes, coronaSPAdes, rnaSPAdes and rnaviralSPAdes version 3.15.3
2. Trinity v2.11.0
3. IVA v1.0.8
4. MEGAHIT v1.2.9
5. PRICE 1.2
6. SAVAGE 0.4.2

### 2 Assembly command lines

#### 2.1 Quality trimming

Prior to assembly all reads underwent quality trimming using BBDuk:

```
bbduk.sh threads=16 trimpolya=15 qtrim=rl trimq=10 <input> <output>
```

#### 2.2 SPAdes, metaSPAdes, coronaSPAdes, rnaSPAdes and rnaviralSPAdes

metaSPAdes, coronaSPAdes, rnaSPAdes and rnaviralSPAdes were run with the default parameters using 16 CPUs, e.g.:

```
coronaspades.py -t 16 -1 left.fastq -2 right.fastq
```

HIV and Influenza HMMs were provided to `rnaviralspades.py` via `--custom_hmms` option:

```
rnaviralspades.py -t 16 -1 left.fastq -2 right.fastq --custom_hmms <hmms>
```

#### 2.3 Trinity

Trinity was run with the default parameters using 16 CPUs and 128 Gb maximum memory:

```
Trinity --seqType fq --left left.fastq --right right.fastq --CPU 16 --max_memory 128G
```

#### 2.4 MEGAHIT

MEGAHIT was run with the default parameters using 16 CPUs and utilizing 10% of available memory:

```
megahit -1 left.fastq -2 right.fastq -t 16 -m 0.1
```

#### 2.5 IVA

IVA was run with the default parameters using 16 CPUs:

```
iva -v --threads 16 -f left.fastq -r right.fastq
```

#### 2.6 PRICE

PRICE assembler was run with parameters suggested by its author for novel viral assembly using 16 threads:

```
PriceTI -fpp left.fastq right.fastq 600 90 -icf seed.fa 1 1 1
-nc 10 -dbmax 151 -mol 30 -tol 20 -mpi 80 -target 90 0 -trimB 0 1.5 -a 16
```

#### 2.7 SAVAGE

The `patch_num` parameter of SAVAGE was chosen, as recommended by SAVAGE authors, to ensure that $500 < read\_coverage/patch\_num < 1000$. We used average viral read coverage as `read_coverage` parameter. SAVAGE was run using 16 threads:

```
savage --split patch_num --p1 left.fastq --p2 right.fastq -t 16
```

### 3 Running time and memory consumption benchmarking

All assemblers were launched in 16 threads on a server with 1.5 TB of RAM and 128 Intel Xeon E7-4880 v2 2.5 GHz cores. Results for SAVAGE are absent as it did not finish after 1 week of runtime.

| | Fr4nk | | Ginger | | PEDV | |
|---|---|---|---|---|---|---|
| | Running time | Memory consumption | Running time | Memory consumption | Running time | Memory consumption |
| MEGAHIT | 1h 23m | 4.3 GB | 1h 9m | 6.9 GB | 1h 46m | 7.3 GB |
| metaSPAdes | 10h 15m | 94.7 GB | 3h 36m | 28 GB | 5h 23m | 33.7 GB |
| rnaSPAdes | 1h 25 m | 16.2 GB | 1h 1m | 16.3 GB | 2h 5m | 22.3 GB |
| Trinity | 19h 32m | 128.0 GB | 17h 42m | 128.0 GB | 23h 5m | 128.0 GB |
| coronaSPAdes | 1h 27m | 14.9 GB | 1h 23m | 13.7 GB | 2h 3m | 19.7 GB |
| rnaviralSPAdes | 1h 23 m | 14.6 GB | 1h 10m | 12.3 GB | 1h 57m | 19.2GB |
| IVA | 2d 5h 46m | 1.5 GB | 3d 10h 17m | 2.1 GB | 5d 3h 8m | 3.7 GB |
| PRICE | 1h 45m | 2.7 GB | 2h 30m | 2.3 GB | 5h 12m | 3.7 GB |
| SAVAGE | N/A | N/A | N/A | N/A | N/A | N/A |

Table 1: Running times and memory consumption of assemblers on coronaviral datasets

### 4 Serratus results

coronaSPAdes was used in the Serratus project for a widespread search of novel CoV and CoV-like species from public sequencing libraries. From a screen of 3.8 million public RNA-seq, meta-genome, meta-virome and meta-transcriptome datasets deposited in NCBI SRA comprising 5.6 petabases of sequencing reads, 52,772 runs potentially containing CoV sequencing reads were identified. 11,120 of the resulting assemblies contained putative CoV contigs, of which 4,179 aligned to CoV RdRp. These assemblies include sequences from 13 previously uncharacterized or unavailable CoV or CoV-like operational taxonomic units (OTUs), defined by clustering amino sequences of the RdRp gene at 97% identity. 8 of these OTUs were designated to a putative novel genus of coronaviruses, noting that all were found in samples from non-mammal aquatic vertebrates falling outside deltacoronaviruses genus.

The length distribution for assemblies of SRA datasets classified as likely CoV-positive, showing a peak around the typical CoV genome length is presented on Figure 1.

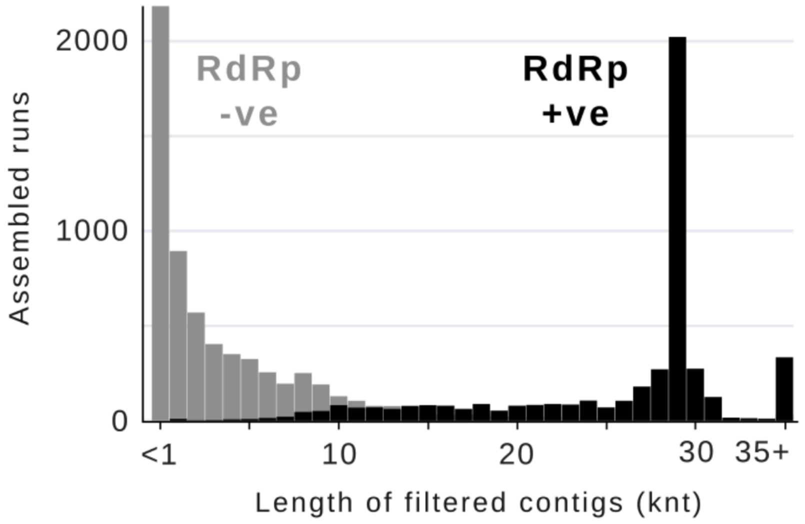


Figure 1: Length distribution for 11,120 assembled contigs classified as CoV-positive, showing a peak around the typical CoV genome length; 4,179 (37.58%) of contigs also contained a match for RdRP.
